# Estimating greater sage-grouse population sizes within the state of Oregon, USA 2017–2024

**DOI:** 10.1101/2025.08.01.668228

**Authors:** Brian G. Prochazka, Peter S. Coates, Austin L. Nash, Shawn T. O’Neil, Skyler T. Vold, Adrian P. Monroe, Cameron L. Aldridge

**Affiliations:** U.S. Geological Survey, Western Ecological Research Center, Dixon Field Station, 800 Business Park Drive, Suite D, Dixon, CA, USA, 95620-9648; Oregon Department of Fish and Wildlife, 237 Highway 20 South, P.O. Box 8, Hines, OR, USA, 97738-9428; U.S. Geological Survey, Fort Collins Science Center, 2150 Centre Avenue, Building C, Fort Collins, CO, USA, 80526-8118

**Keywords:** abundance, lek attendance, N-mixture model, population size, sex-ratio, simulation

## Abstract

We fit an *N*-mixture model to lek (breeding area) count data to estimate annual population sizes of greater sage-grouse (*Centrocercus urophasianus*; sage-grouse) within the state of Oregon, USA between 2017–2024. Population estimates were delineated among 24 Priority Areas for Conservation (PACs) and considered additional sources of information including male-to-female sex ratios, lek attendance rates, numbers of unmodeled leks, and the existence of unsampled/unknown leks. In 2024, the state of Oregon was estimated to contain approximately 41,875 sage-grouse (95% credible interval [CRI] = 38,980–54,634), which was down from a high of 50,869 (95% CRI = 41,794–66,238) in 2017. A nadir (low point) was identified during 2019, when the median statewide population estimate was 30,644 birds. A complete population oscillation was not evident during the inferential period based on local maxima that were observed during the start (2017) and stop (2024) years of analysis. In addition to estimating population sizes, we evaluated *N*-mixture model estimates for precision and accuracy after randomly removing single and repeat counts in 10% increments (relative to total sample size). We estimated an increase in absolute bias of approximately 1.6% for every 10% reduction in effort.

## Introduction

Estimating the size and trajectory (trend) of wildlife populations is a key component of effective species management. For greater sage-grouse (*Centrocercus urophasianus*; hereafter, sage-grouse), there has been a wealth of information about trends, but considerably less has been published on population size. This is particularly troubling considering the prevalence and duration of population declines documented throughout much of the species’ range (Coates et al., 2021). The few studies that have presented information on population size have been relatively narrow in geographic scope, presented the value as an index thereby limiting its utility, or failed to account for leks that were not modeled due to data limitations or logistical constraints (Garton et al., 2011; McCaffery and Lukacs, 2016; Shyvers et al., 2018; Coates et al., 2021). Historically popular approaches to modeling population size have required data that are difficult to collect at scale (e.g., time intensive), such as individual identification (e.g., capture–recapture) or systematic repeated counts that use distance sampling to estimate detectability. Sage-grouse are a cryptic, low-density species when not on their breeding sites (i.e., leks), so commonly used bird monitoring protocols (e.g., Integrated Monitoring in Bird Conservation Regions, Breeding Bird Survey, eBird) are inadequate because detections are too few and far between. This, in part, could explain the current lack of information on population sizes for sage-grouse across their geographic range and through time.

Recent advances in population modeling have provided an opportunity to better address questions of population size by estimating a biologically meaningful value (“true” abundance) from data that can be collected over large spatiotemporal scales with relative ease. Using only replicated counts of individuals across space and time, these models (hereafter, N-mixture models) can produce separate estimates of latent abundance and detection probability. Covariates can be fit within state (i.e., latent abundance) or observation likelihoods to improve parameter estimation and inform future sampling designs (e.g., when or how intensively to focus monitoring efforts). Additional sources of information can be included, within or outside the model, to further refine abundance estimates. Specific to sage-grouse, these additional sources of information could include male-to-female ratios, lek attendance rates, and numbers of unknown or unmodeled active leks. Here, we estimated sage-grouse population abundance in Oregon, USA, using N-mixture models of sage-grouse male counts at leks, combined with ancillary harvest data to inform sex ratios, priors on lek attendance to account for the non-breeding or non-lek-breeding cohort, derived parameters to account for unmodeled leks, and complementary statistical methods to estimate currently undetected leks. Results improve upon the previous best available estimates on population size for the state of Oregon and can be used directly in conservation and land management plans within the state.

## Methods

### Data

We used sage-grouse lek count data provided by the Oregon Department of Fish and Wildlife (ODFW) to estimate annual population sizes of sage-grouse within the state of Oregon between 2017–2024. Data spanned the geographic range of sage-grouse within the state (Fig. 1), with all lek counts spatially linked to one of 24 population units referred to as Priority Areas for Conservation (PACs; included Low-Density areas). Leks located outside of PAC boundaries were assigned to the nearest PAC using the sf package (v 1.0-16) in R (R Core Team, 2024).

**Figure 1.**
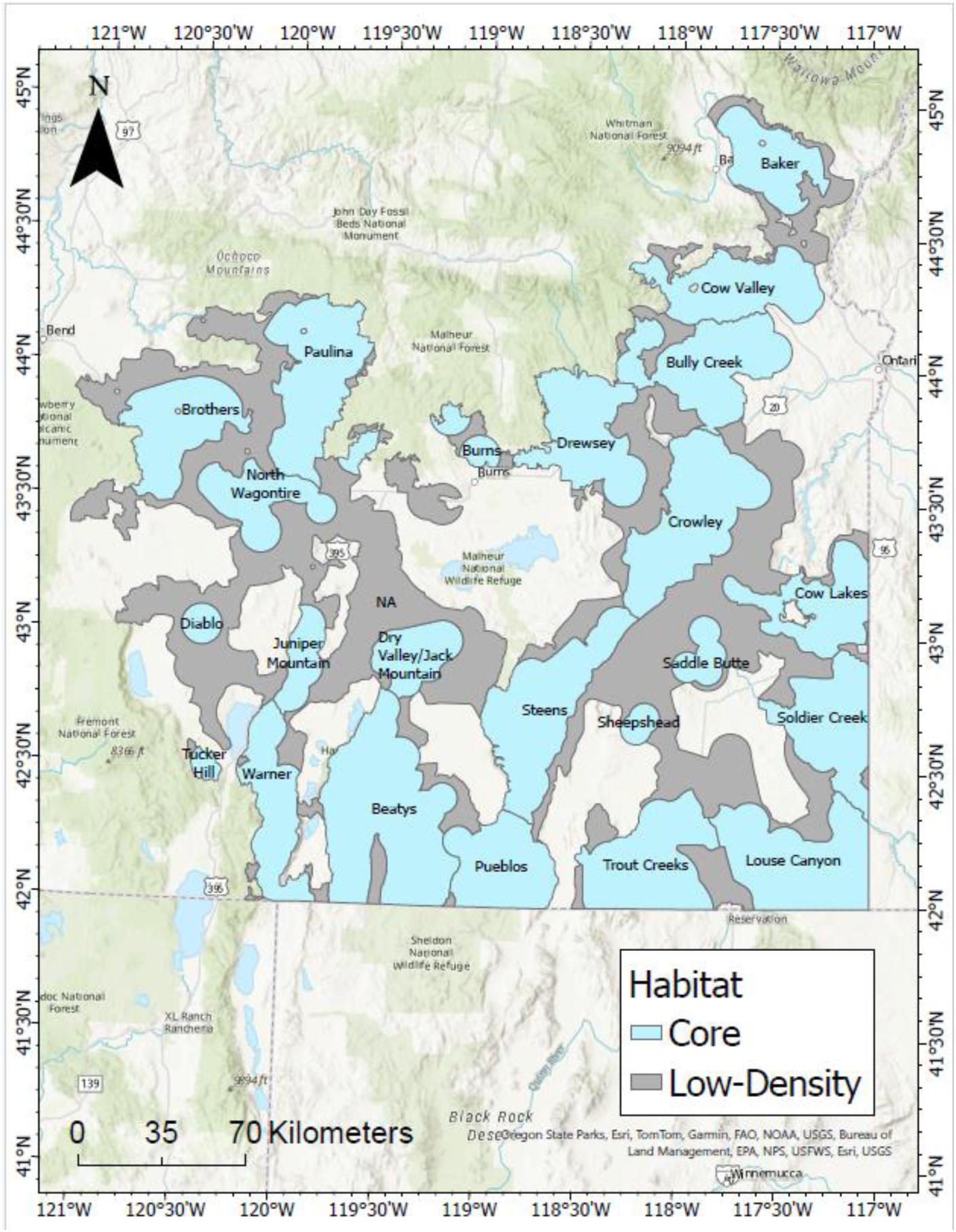
Greater sage-grouse (*Centrocercus urophasianus*) Priority Areas for Conservation within the state of Oregon, USA. Habitat is delineated as Core (cyan polygons) and Low-Density (gray polygons).

To produce a dataset appropriate for modeling, we first filtered lek count data using previously established methods (Monroe et al., 2016; O’Donnell et al., 2021). We selected counts that were conducted within seasonal and diurnal periods of peak lek attendance. Seasonal peaks in lek attendance were targeted by restricting the dataset to counts occurring between March 1 and May 31 of each year (Blomberg et al., 2013; Jenni and Hartzler, 1978; Wann et al., 2019). Diurnal peaks in lek attendance were targeted by restricting the dataset to counts occurring between 30 minutes before and 120 minutes after local sunrise (Monroe et al., 2016). When multiple counts were recorded during the same day, we retained the maximum count, assuming the count met date and time criteria. We also required leks to be counted at least once per year during five separate years (between 2017–2024) with at least two years producing a maximum count that was ≥2 males.

### Investigating rates of lek switching

Prior to modeling, we investigated the potential for high rates of lek switching (i.e., individual male sage-grouse attending >1 lek within the same breeding season) among proximal leks, which if left unresolved could yield double counts and inferences about population size that were biased high. We therefore constructed two separate matrices from the filtered lek count dataset. The first was a distance matrix representing pairwise distances among all leks. The second was a time series correlation matrix of Pearson’s correlation coefficients constructed from pairwise combinations of coincident lek counts (lek counts conducted on the same day). Lek pairs with fewer than five coincident counts were discarded from this portion of the analysis. Correlation coefficients were then grouped according to pairwise distances binned at 200-m increments (range = 0–374.6 km). Significant rates of lek switching were evidenced by distributions of correlation coefficients that were entirely below zero. Approximately 2.6% of the 1,793 distance bins investigated had a maximum correlation coefficient less than zero. The only proximal bin to have a maximum correlation coefficient less than zero was the >0–200-m bin (mean = -0.25; range = -0.41 – -0.11). The remaining bins with a maximum correlation coefficient less than zero were from leks that were >315-km apart, which precluded a lek switching mechanism. Due to the small number of leks (<1%) that could be affected by lek switching and the relatively low rate (evidenced by a moderately negative correlation coefficient), we opted to model all leks independently rather than aggregate proximal leks.

### Identifying active, unmodeled leks

We recorded the number of leks that did not meet filtering criteria and were therefore not included in models. Of those unmodeled leks, we identified a subset that provided some evidence of continued activity (≥2 counts of ≥1 male between 2017–2024 and no zero counts during 2023–2024) and used them to calculate a derived abundance parameter in our model (detailed in modeling section). We refer to those unmodeled active leks as *leks*^+^. For the leks that were modeled, we converted the count date and time information to ordinal date and time since sunrise (TSSR). Ordinal date and TSSR were standardized by subtracting the sample mean and dividing by the sample standard deviation.

### Modeling lek count data

We fit an *N*-mixture model (Kéry and Schaub, 2012; Royle, 2004) to repeated counts, across space and time, of male sage-grouse attending leks. This yielded separate estimates of latent abundance and detection probability. The latent abundance (𝑁_𝑙𝑡_) for each lek (𝑙) and year (𝑡) were modeled using a state-space framework, such that 𝑁_𝑙𝑡_ was defined as the product of the previous year’s abundance (𝑁_𝑙𝑡−1_) and finite rate of population change (𝜆_𝑙𝑡−1_):

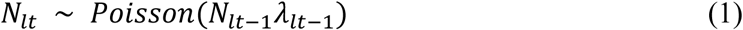

Initial population sizes were specified using uniformly distributed priors bounded by half the minimum male count (lower bound) and twice the maximum male count (upper bound) observed per lek between 2017–2024:

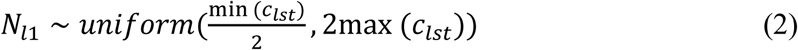

Lek-level estimates of population change were modeled on a log-scale using a normal distribution and vague priors for mean and standard deviation parameters:

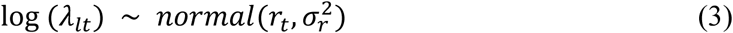

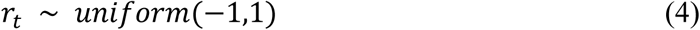

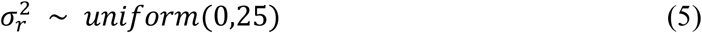

The observed counts (𝑐_𝑙𝑠𝑡_), which included within-year temporal replicates (𝑠), were assumed to follow a binomial distribution with count-level detection probability (𝑝_𝑙𝑠𝑡_):

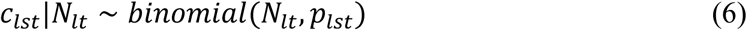

Covariates for ordinal date and TSSR were modeled on the logit scale, allowing for the mean detection probability to vary across a season and within a morning (Monroe et al., 2019):

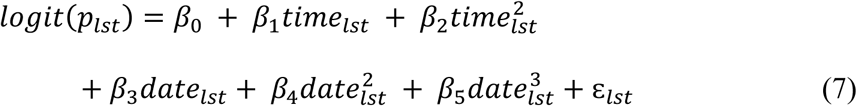

The error term in our model (ɛ_𝑙𝑠𝑡_) was normally distributed with mean 0 and standard deviation

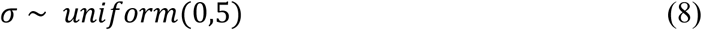

### Simulations to evaluate sampling effort

To explore the influence of sampling effort on estimation of abundance, we ran four separate models on progressively smaller, subsampled versions of the dataset by randomly reassigning observed counts with NA values. These simulated levels of varying sampling intensity resulted in distributional shifts in the proportion of leks receiving 0–3 counts within a single breeding season (Fig. 2).

**Figure 2.**
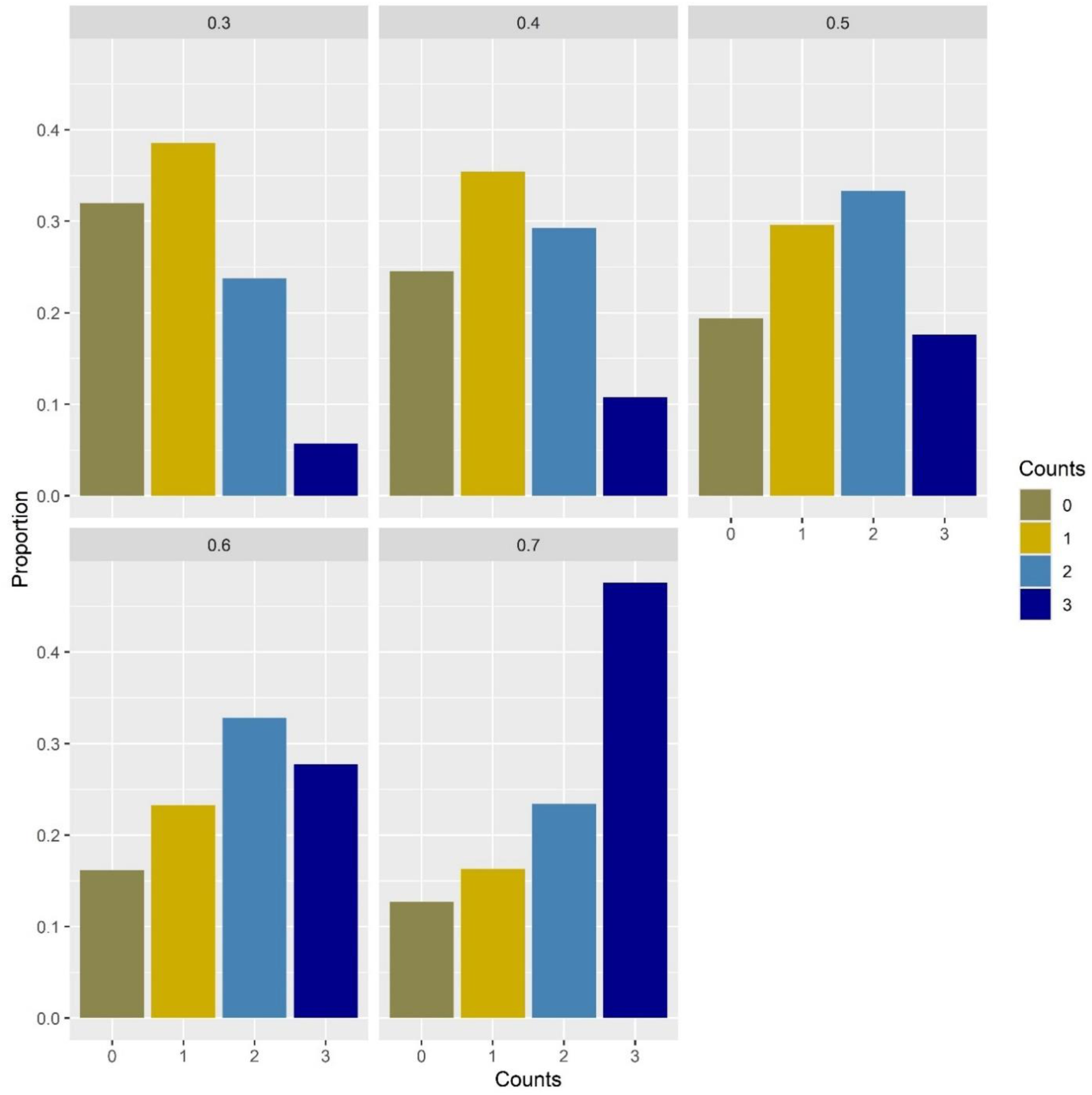
The average proportion of greater sage-grouse (*Centrocercus urophasianus*) leks (breeding areas) that received 0–3 counts per breeding season (recorded during separate days) between 2017–2024. Individual panels depict varying levels of lek count effort with 30–70% (listed in title; 0.3–0.7) of leks receiving 2–3 counts per breeding season (i.e., repeat counts; blue tones). The reference dataset (0.7) depicts current maximal lek count efforts carried out by the Oregon Department of Fish and Wildlife, and their partners, based on data that were deemed suitable for *N*-mixture model analysis (see Methods for more detailed information on lek count criteria). Proportions less than 0.7 represent smaller datasets created from random, simulated reductions in the number of observed counts.

Lek count efforts during the most recent 8-year period (2017–2024) resulted in repeat counts (2– 3 counts per breeding season) for approximately 70% of modeled leks. Simulated reductions in sampling effort resulted in 30–60% of modeled leks having within-season repeat counts (*reduced models*). Parameter estimates of abundance from *reduced models* were compared to estimates from the same model fit to data collected under the current maximal sampling intensity (i.e., 70%, *reference model*) to evaluate the impacts of increasingly sparse data on parameter accuracy and precision. We calculated bias and coverage statistics for the derived parameters of abundance calculated at the PAC level. Bias was calculated as the mean difference between the median estimate of the *reference model* and the median estimate of the *reduced model*, divided by the median estimate of the *reference model* (multiplied by 100 to produce a percentage). Coverage was calculated as the percent of the *reduced model’s* 90% credible intervals (CRIs) that overlapped the median of the *reference model’s* counterpart parameters.

### Derived parameters for improved estimation of abundance

Despite advantages of working within an *N*-mixture framework, lek count data still impose several limitations on the scope of model inference. First, lek count data are typically restricted to males and even when females are included, the sex ratio may not reflect the true population value. To obtain estimates of total population size, additional information about the female proportion of the population must be included. Estimates of male population size can be supplemented by assuming a regionally constant sex ratio. However, previous studies conducted from different areas of the geographic range of sage-grouse have provided evidence for spatial variation in sex ratios (Guttery et al., 2013; Prochazka et al., 2024; Shyvers et al., 2023). To obtain a sex ratio estimate specific to Oregon, we utilized hunter-harvested wing data collected by the ODFW between 1995–2023, corrected for age-specific harvest susceptibility (Hagen et al., 2018). The resultant, long-term mean sex ratio of 1.82 females per male (95% CRI = 1.76– 1.88) was multiplied by *N*-mixture estimates of male abundance to obtain an estimate of female abundance.

Another limitation inherent to lek count data is that not all males attend at least one lek within a breeding season (Blomberg et al., 2013; Wann et al., 2019). This is especially true for juvenile or yearling males, which may utilize areas proximal to leks, but still outside of detectable range. Because these individuals are not available for detection, the model cannot account for their absence and additional information is required to account for their contribution to total population size (Monroe et al., 2019). For this, we used an informative prior from a study of GPS-marked male sage-grouse monitored during multiple breeding seasons (Wann et al., 2019), and which represented the age-weighted average peak lek attendance rate (0.58; 95% CRI = 0.46–0.71).

The final issue with lek count data, as it pertains to estimates of total population size, is that not all active leks in existence are present in the data. To account for this, we estimated the number of unknown leks likely to exist on the landscape at present using a larger subset of the ODFW lek count database that spanned 85 years (1940–2024). First, we calculated the number of new leks added to the database on an annual basis between 1940–2023. This calculation was conducted at the PAC level using a cumulative sum for each year. For example, the number of unknown leks per PAC in 2022 was calculated as the sum of new leks added per PAC during 2023 and 2024. We then used an autoregressive technique (Generalized Regression Neural Network [GRNN]; *tsfgrnn* package, v 1.0.5) to forecast a time series of future values (Martínez et al., 2022) per PAC and used the value at time step 𝑡 = 1 (i.e., number of new leks likely to be discovered beyond 2024) to represent the number of unknown leks likely to exist at present time (Fig. 3; Table 1).

**Figure 3.**
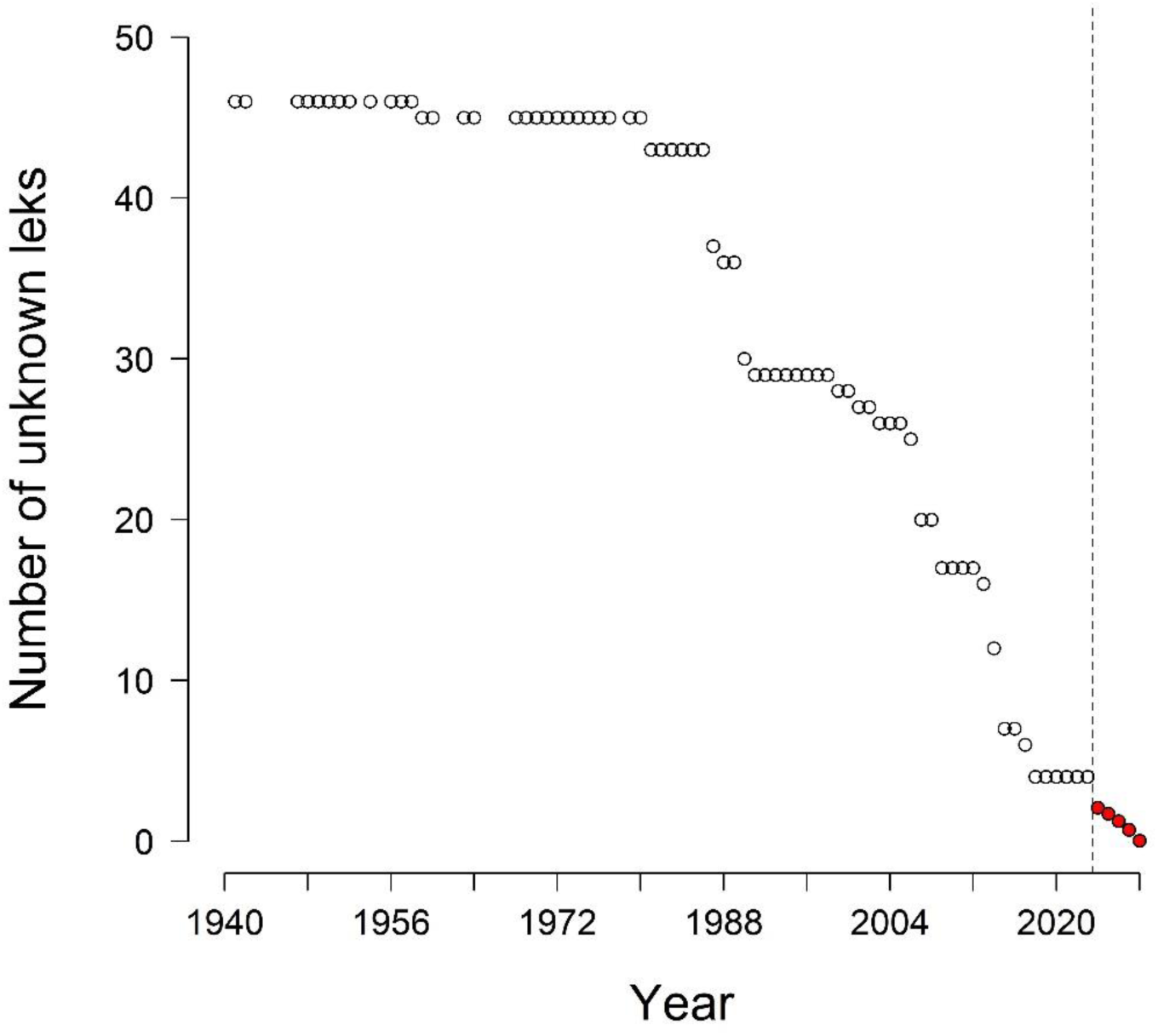
Number of greater sage-grouse (*Centrocercus urophasianus*) leks (breeding areas) missing from the Oregon Department of Fish and Wildlife (ODFW) lek count database between 1940–2024, within the Pueblos Priority Area for Conservation (PAC), calculated as the cumulative sum of new leks added to the database during all subsequent years. Predictions (red circles) of the number of unknown leks likely to exist during four future years, based on the application of a Generalized Regression Neural Network (GRNN) to the observed data (open circles).

**Table 1.** Predictions of the number of unknown greater sage-grouse (*Centrocercus urophasianus*) leks (breeding areas) likely to exist (during 2024) within Priority Areas for Conservation (PACs) based on the application of a Generalized Regression Neural Network (GRNN) to lek count data collected within the state of Oregon, USA between 1940–2024.

| Priority Areas for Conservation | Estimated number of unknown leks |
| --- | --- |
| Baker | 0 |
| Beatys | 7 |
| Brothers | 0 |
| Bully Creek | 0 |
| Burns | 0 |
| Cow Lakes | 0 |
| Cow Valley | 0 |
| Crowley | 0 |
| Diablo | 0 |
| Drewsey | 1 |
| Dry Valley/Jack Mountain | 1 |
| Juniper Mountain | 0 |
| Louse Canyon | 3 |
| Low-Density Areas | 0 |
| North Wagontire | 0 |
| Paulina | 0 |
| Pueblos | 3 |
| Saddle Butte | 0 |
| Sheepshead | 0 |
| Soldier Creek | 3 |
| Steens | 3 |
| Trout Creeks | 0 |
| Tucker Hill | 0 |
| Warner | 0 |

Each of the previous corrections applied to *N*-mixture estimates were based on inherent limitations of lek count data. Limitations that we introduced prior to modeling, in the form of observation criteria (e.g., date of count, time of count, number of counts), resulted in an even narrower geographic scope. We corrected for these user-imposed limitations by calculating derived parameters of abundance based on PAC-level numbers of unmodeled leks (i.e., *leks*^+^). Like the GRNN predictions of numbers of unknown leks (i.e., missing from database altogether), we multiplied the annual average, PAC-specific, lek-level abundance (corrected for attendance and sex-ratios) by the number of unmodeled, active leks within a PAC.

### Model tuning and investigation

All models were run using Markov chain Monte Carlo techniques implemented in JAGS (Plummer, 2003) using the statistical programming language and software package R (R Core Teams, 2024). We ran 150,000 iterations on each of three chains, discarding the initial 100,000 iterations as burn-in. The remaining 50,000 iterations were thinned by a factor of 50, resulting in 3,000 posterior samples for inference. For each model we evaluated fit using posterior predictive checks (Bayesian p-values) based on chi-square discrepancy test statistics (Conn et al., 2018; Kéry and Royle, 2015). We assessed model convergence using the r-hat statistic (Brooks and Gelman, 1998).

## Results

The ODFW lek database, restricted to the years 2017–2024, contained 13,521 individual lek counts from 1,182 leks. The average number of counts per lek was 11.44 (SD = 7.78), or 1.43 counts per lek per year (SD = 0.97). When we restricted the dataset to the leks and counts that were modeled (based on a 70% effort for repeated measures) we were left with 6,573 counts recorded from 385 leks. The average number of counts per lek from the reduced dataset was 17.07 (SD = 6.02), which equated to 2.13 counts per lek per year (SD = 0.75). The number of active leks that were not modeled (*leks*^+^) from that dataset was 126. Nearly 40% of active unmodeled leks were in the Beatys (*n* = 29) and Cow Valley (*n* = 19) PACs. Louse Canyon (*n* =12), Trout Creeks (*n* =11), Warner (*n* =10), and Soldier Creek (*n* =10) also had high numbers of active unmodeled leks. The remaining 671 leks, based on our criteria, were either inactive or infrequently counted between 2017–2024.

We found the model to fit the data well based on a Bayesian p-value equal to 0.51 (Fig. 4). The model also demonstrated convergence based on a maximum r-hat statistic of 1.06.

**Figure 4.**
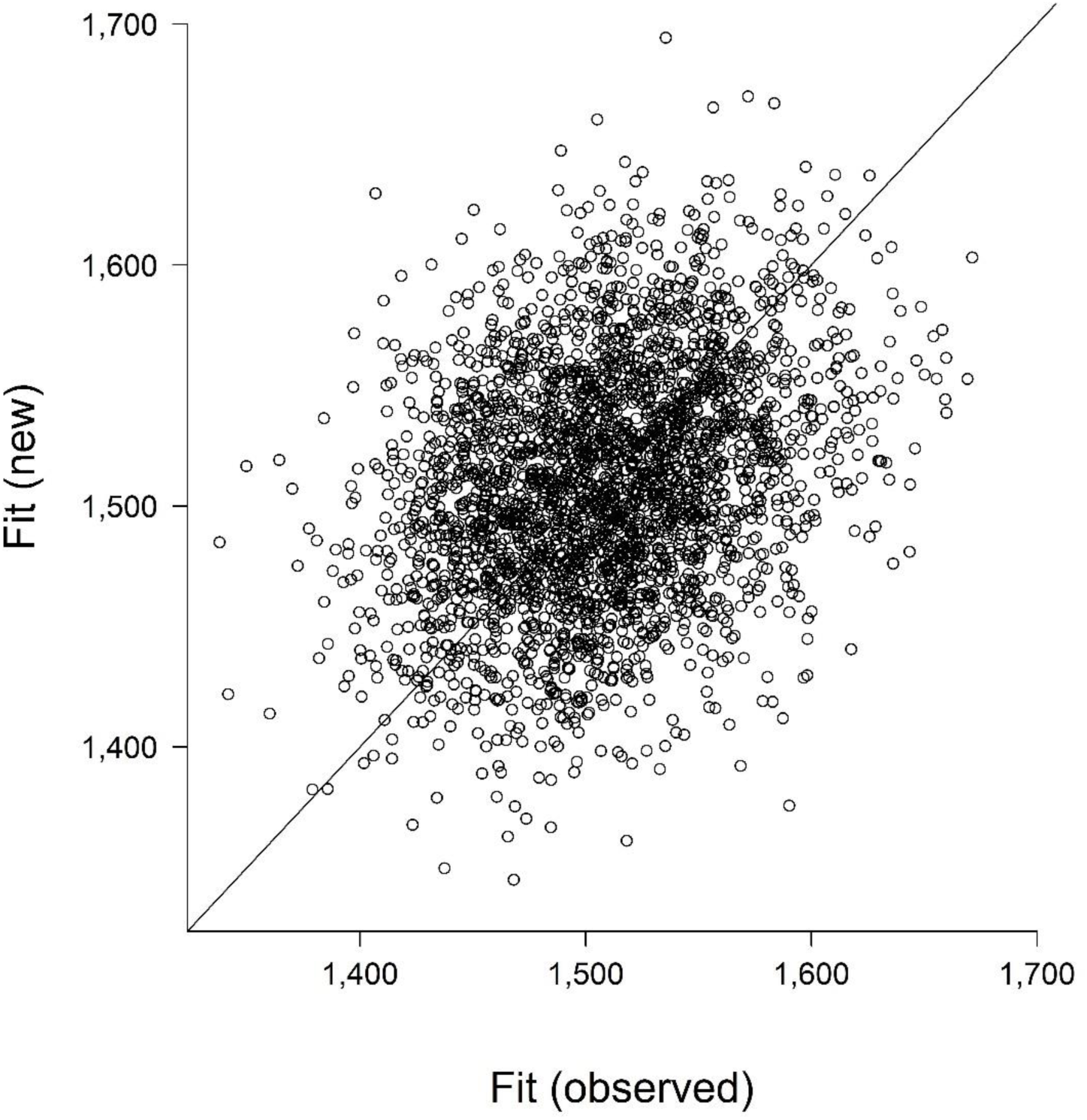
Graphical posterior predictive check of *N*-mixture model adequacy based on chi-square discrepancy measure. The *N*-mixture model was fit to greater sage-grouse (*Centrocercus urophasianus*) lek (breeding area) count data that spanned eight years (2017–2024) and covered the species’ range in the state of Oregon, USA. The Bayesian *p*-value is equal to the proportion of points above the 1:1 line.

The average detection probability across all counts was estimated to be 0.54; however, there was considerable variation (Fig. 5) with the 95% CRI spanning 0.23–0.82.

**Figure 5.**
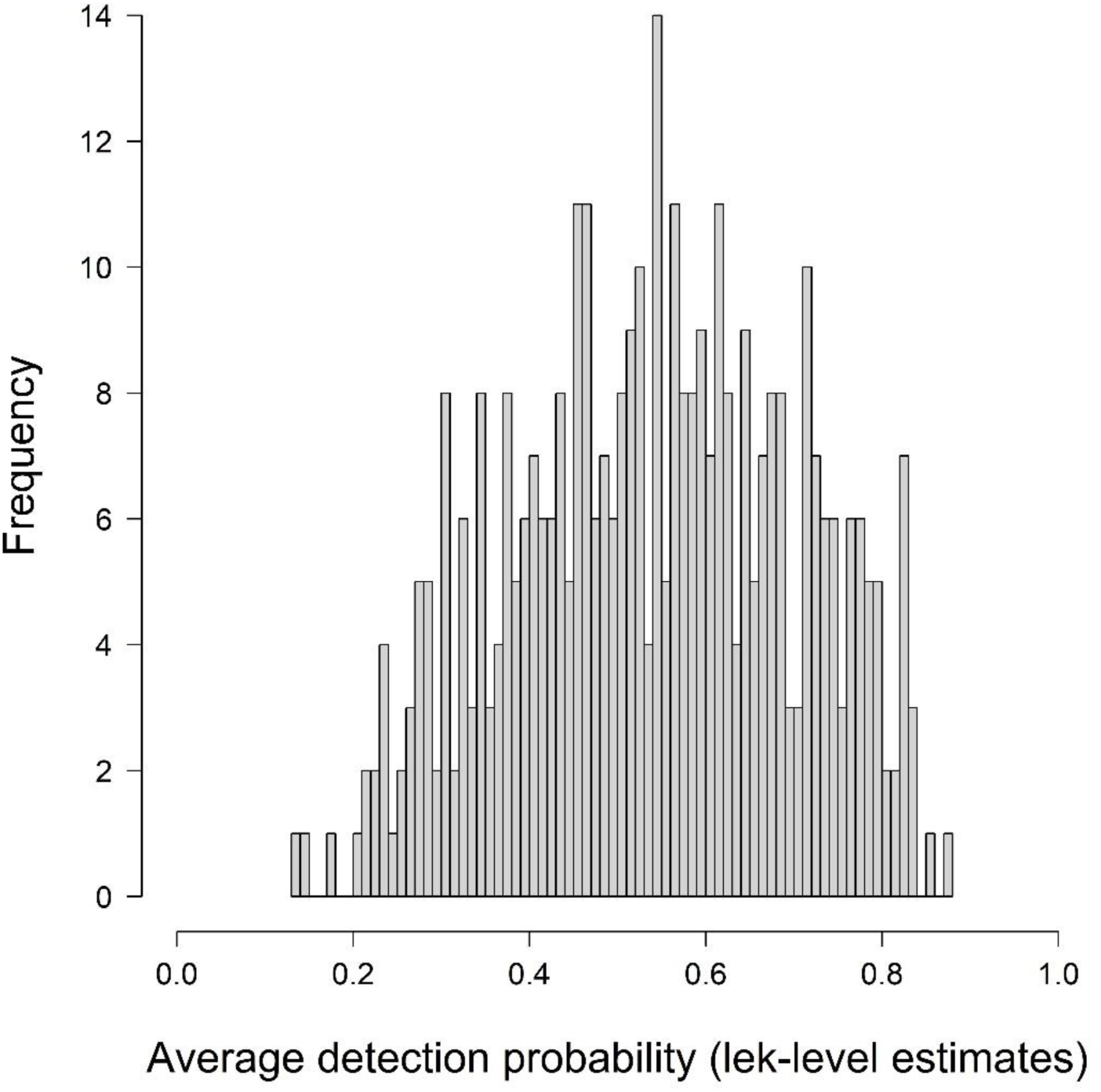
Estimates of the average detection probability of male greater sage-grouse (*Centrocercus urophasianus*) attending leks (breeding areas) based on an *N*-mixture model that was fit to count data spanning eight years (2017–2024) and covering the species’ range in the state of Oregon, USA.

Time since sunrise (TSSR) had a considerable negative effect on detection probability (Fig. 6) with the highest values observed 30 minutes prior to sunrise (0.65; 95% CRI = 0.56–0.72) and the lowest values at 120 minutes after sunrise (0.19; 95% CRI = 0.09–0.33). Detection probabilities remained above 0.5 until 42 minutes after sunrise then declined after.

**Figure 6.**
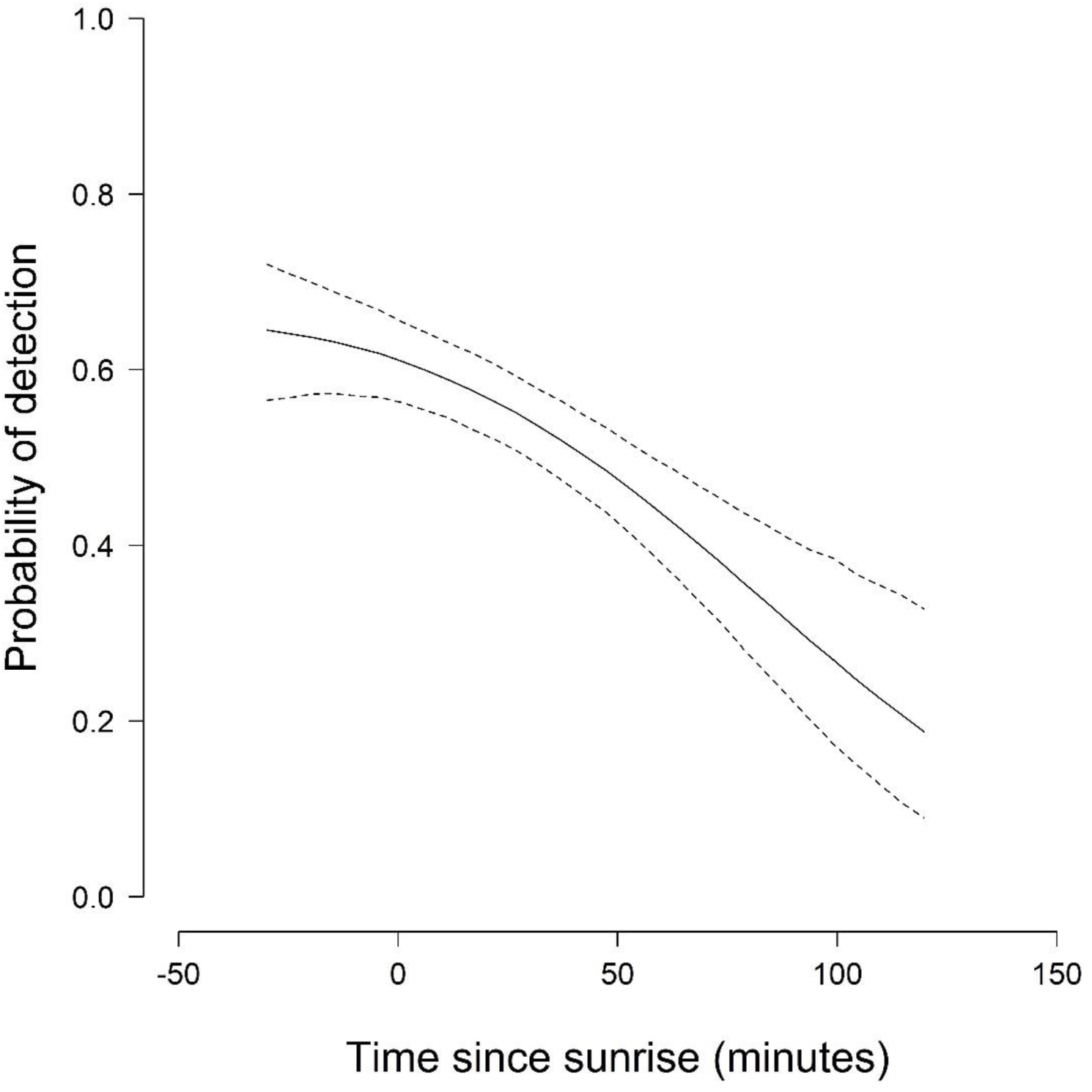
The effect of time since sunrise (in minutes) on the detection probability of male greater sage-grouse (*Centrocercus urophasianus*) attending leks (breeding areas) within the state of Oregon, USA. Predictions are based on an *N*-mixture model fit to lek count data collected between 2017–2024. The median estimate is depicted using a solid black line. The 95% credible interval is depicted using dashed lines.

We did not detect an effect of ordinal date on detection probability (Fig. 7); credible intervals were wide with considerable overlap across all days investigated.

**Figure 7.**
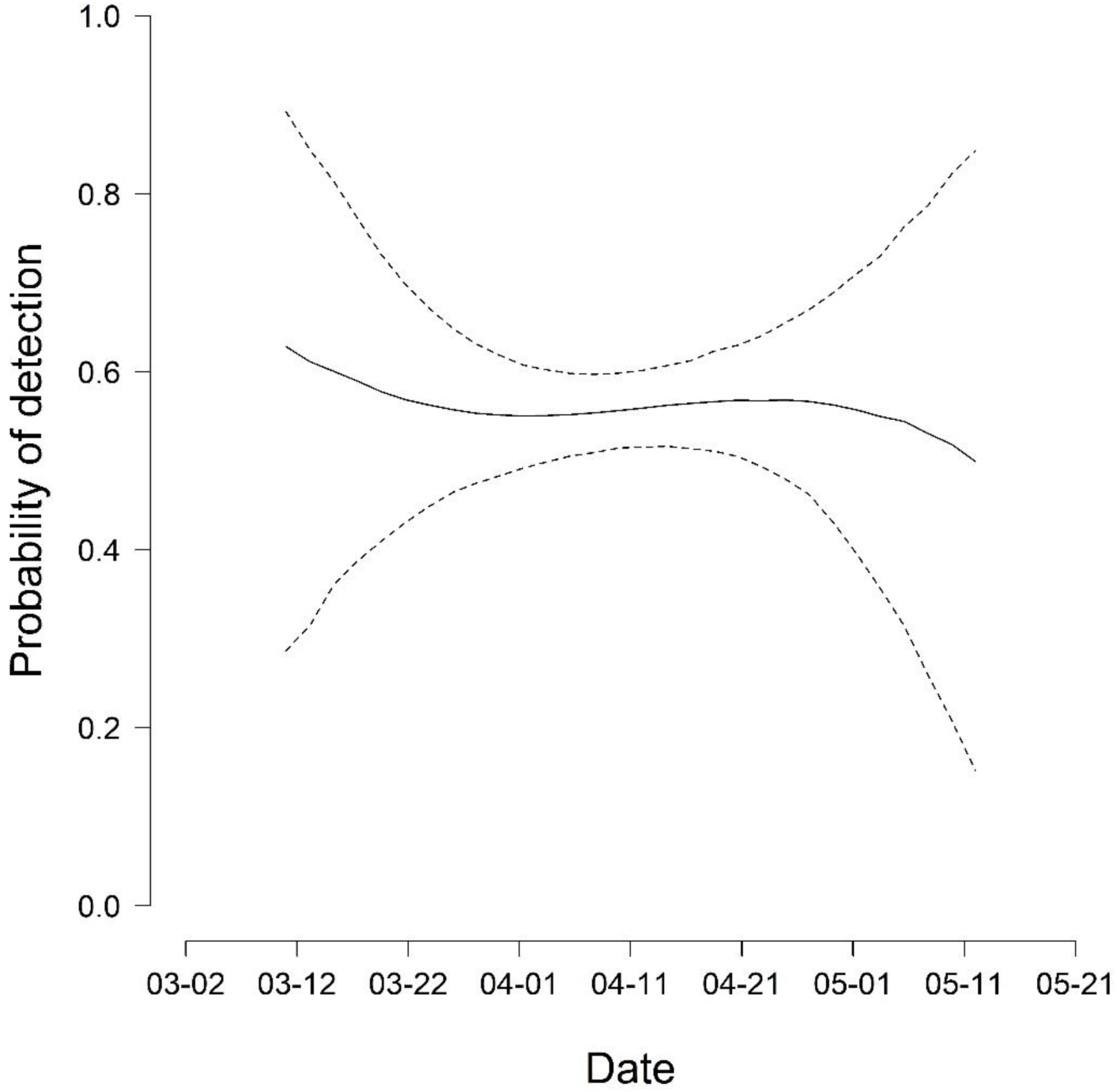
The effect of date on the detection probability of male greater sage-grouse (*Centrocercus urophasianus*) attending leks (breeding areas) within the state of Oregon, USA. Predictions are based on an *N*-mixture model fit to lek count data collected between 2017–2024. The median estimate is depicted using a solid black line. The 95% credible interval is depicted using dashed lines.

In 2024, the state of Oregon was estimated to contain approximately 41,875 sage-grouse (95% CRI = 38,980–54,634), which was down from a high of 50,869 (95% CRI = 41,794–66,238) in 2017. A nadir (low point) was identified during 2019, when the median statewide population estimate was 30,644 birds (95% CRI = 25,106–39,917) (Fig. 8).

**Figure 8.**
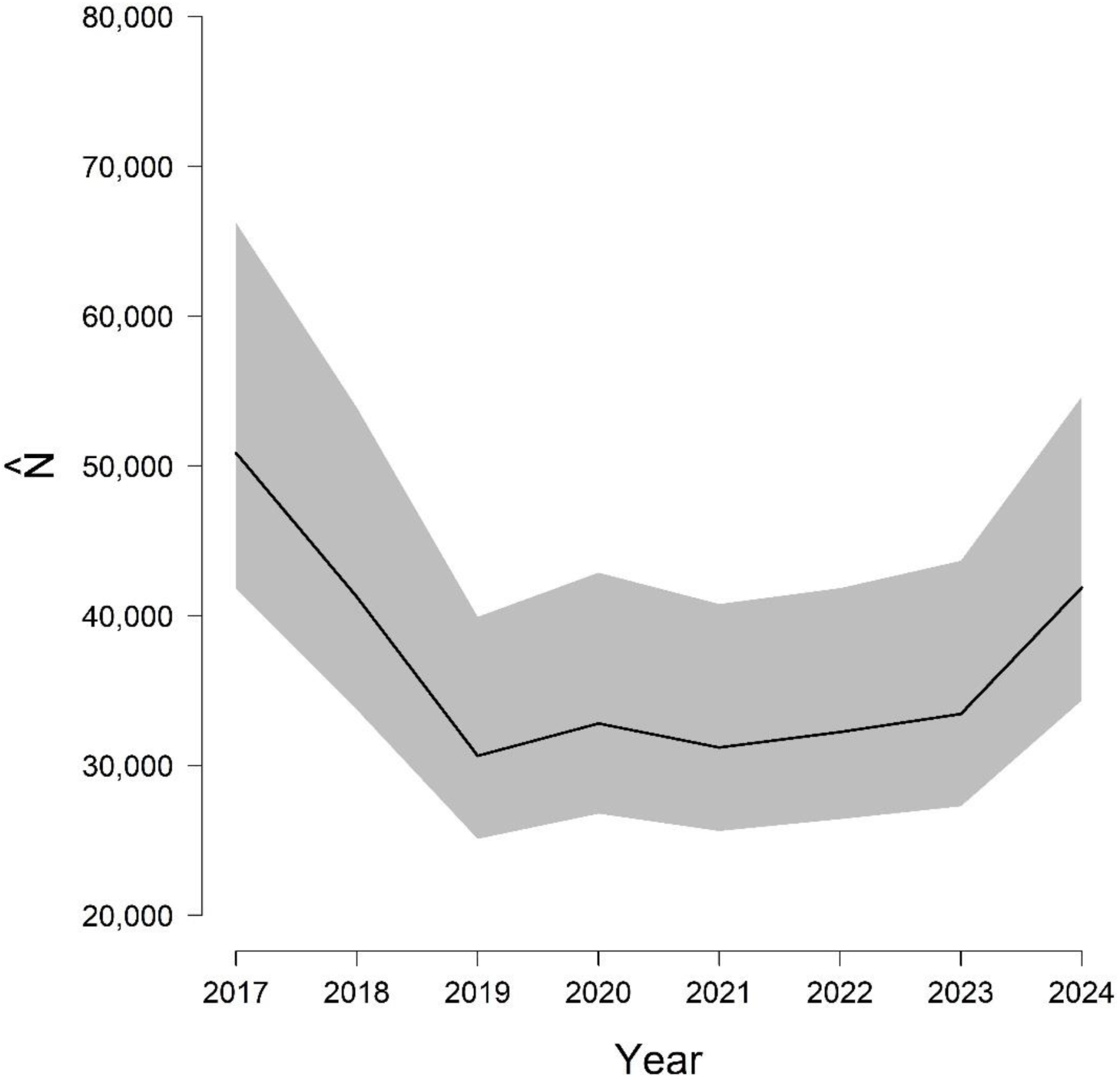
*N*-mixture model estimates of greater sage-grouse (*Centrocercus urophasianus*) abundance for the state of Oregon, USA between 2017–2024. The median (black line) and 95% credible interval (gray polygon) are plotted.

A complete population oscillation was not evident during the inferential period based on local maxima that were observed during the start (2017) and stop (2024) years of analysis. References to population change reported here are likely biased low compared to trends reported elsewhere using nadir-to-nadir trend estimation methods (Coates et al., 2021).

Substantial variation was observed across PACs in both the size and trajectory of the population. The greatest declines were observed within the Cow Lakes, Diablo, North Wagontire, Paulina, Sheepshead, and Soldier Creek PACs (Fig. 9). As of 2024, each of those PACS represented approximately 0.1% (Diablo), 0.4% (Sheepshead), 0.9% (North Wagontire), 2.6% (Cow Lakes), 4.1% (Paulina), and 5.9% (Soldier Creek) of the state’s total population size.

**Figure 9.**
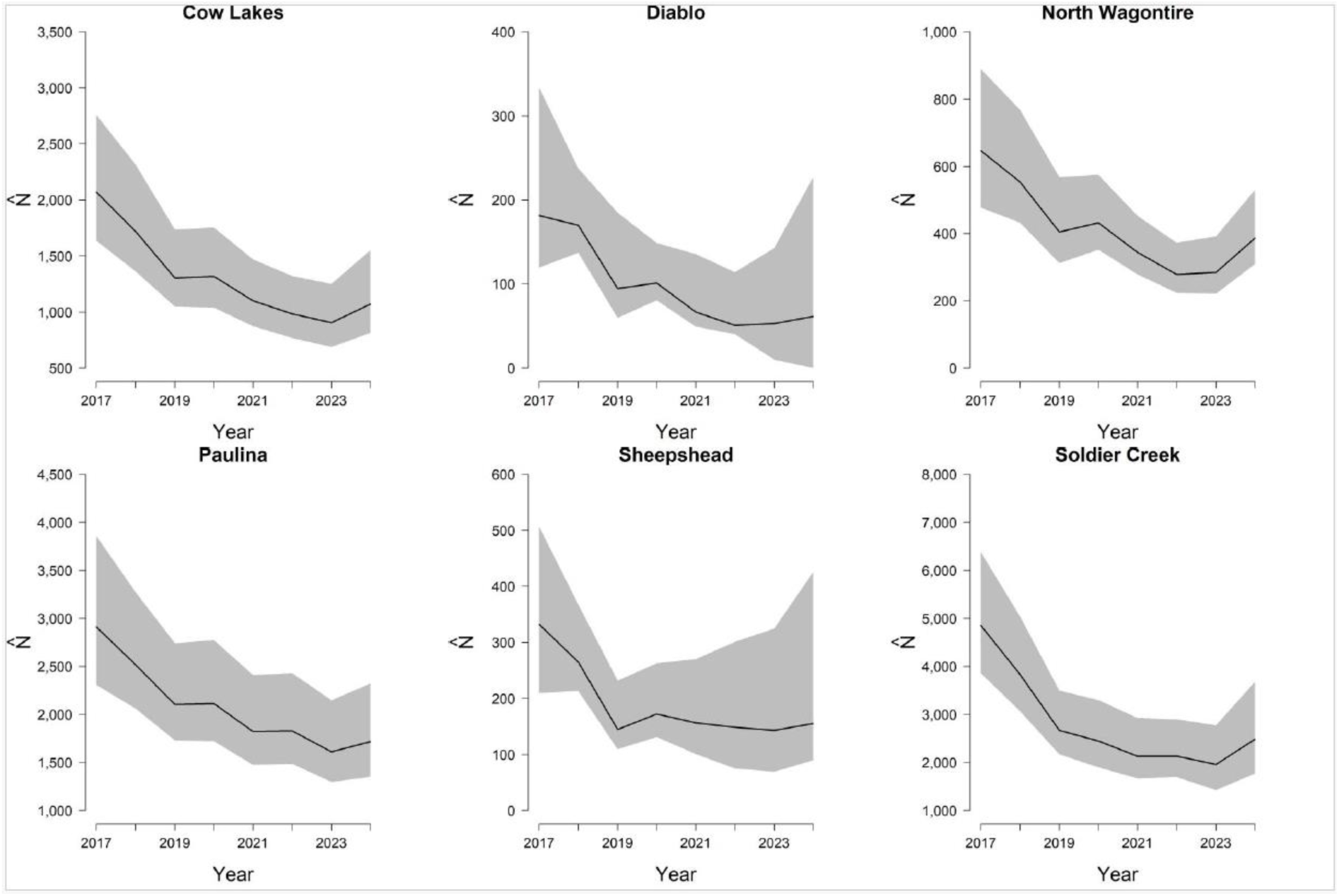
*N*-mixture model estimates of greater sage-grouse (*Centrocercus urophasianus*) abundance within Priority Areas for Conservation (PAC) located in the state of Oregon, USA between 2017–2024. PACs depicted (Cow Lakes, Diablo, North Wagontire, Paulina, Sheepshead, Soldier Creek) are identified in the title of each figure. The median (black line) and 95% credible interval (gray polygon) are plotted for each PAC.

PACs that experienced growth during the 8-year period were Dry Valley/Jack Mountain, Juniper Mountain, Louse Canyon, and Steens (Fig. 10). Those PACs represented 0.6% (Juniper Mountain), 1.8% (Dry Valley/Jack Mountain), 6.7% (Steens), and 11.3% (Louse Canyon) of the state’s total population size during 2024.

**Figure 10.**
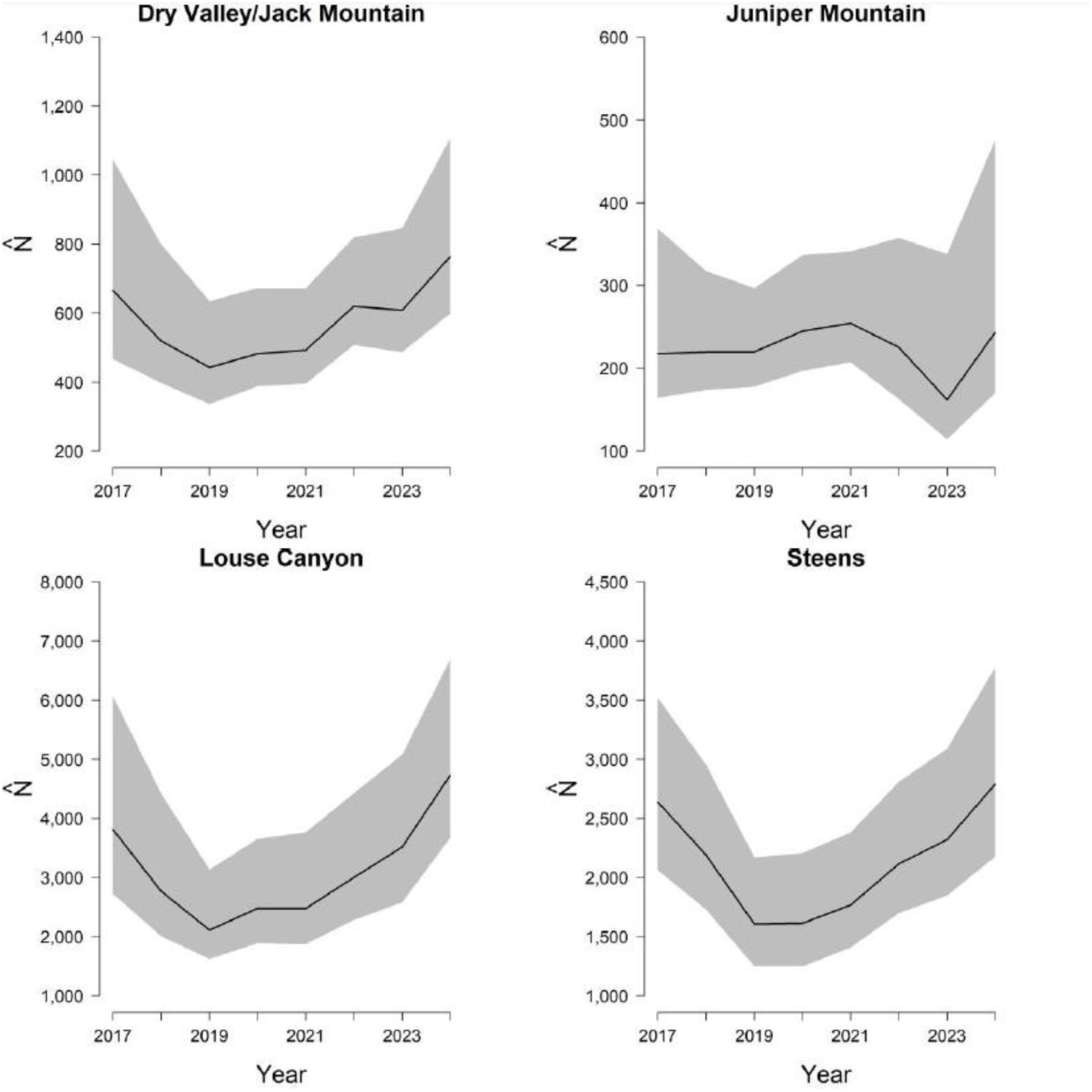
*N*-mixture model estimates of greater sage-grouse (*Centrocercus urophasianus*) abundance within Priority Areas for Conservation (PAC) located in the state of Oregon, USA between 2017–2024. PACs depicted (Dry Valley/Jack Mountain, Juniper Mountain, Louse Canyon, Steens) are identified in the title of each figure. The median (black line) and 95% credible interval (gray polygon) are plotted for each PAC.

Beatys remained the largest PAC across the 8-year period (Fig. 11), losing approximately 7% of its maximum population size (2017 value) and currently sustaining around 7,427 (95% CRI = 5,991–9,846) birds (approximately 17.7% of the 2024 state total). Diablo remained the smallest PAC across the entire timeframe (Fig. 11) and the PAC with the greatest losses overall (approximately 66%). Diablo was estimated to have had 181 birds (95% CRI = 120–334) in 2017 and is now estimated at 61 birds (95% CRI = 0–227) in 2024.

**Figure 11.**
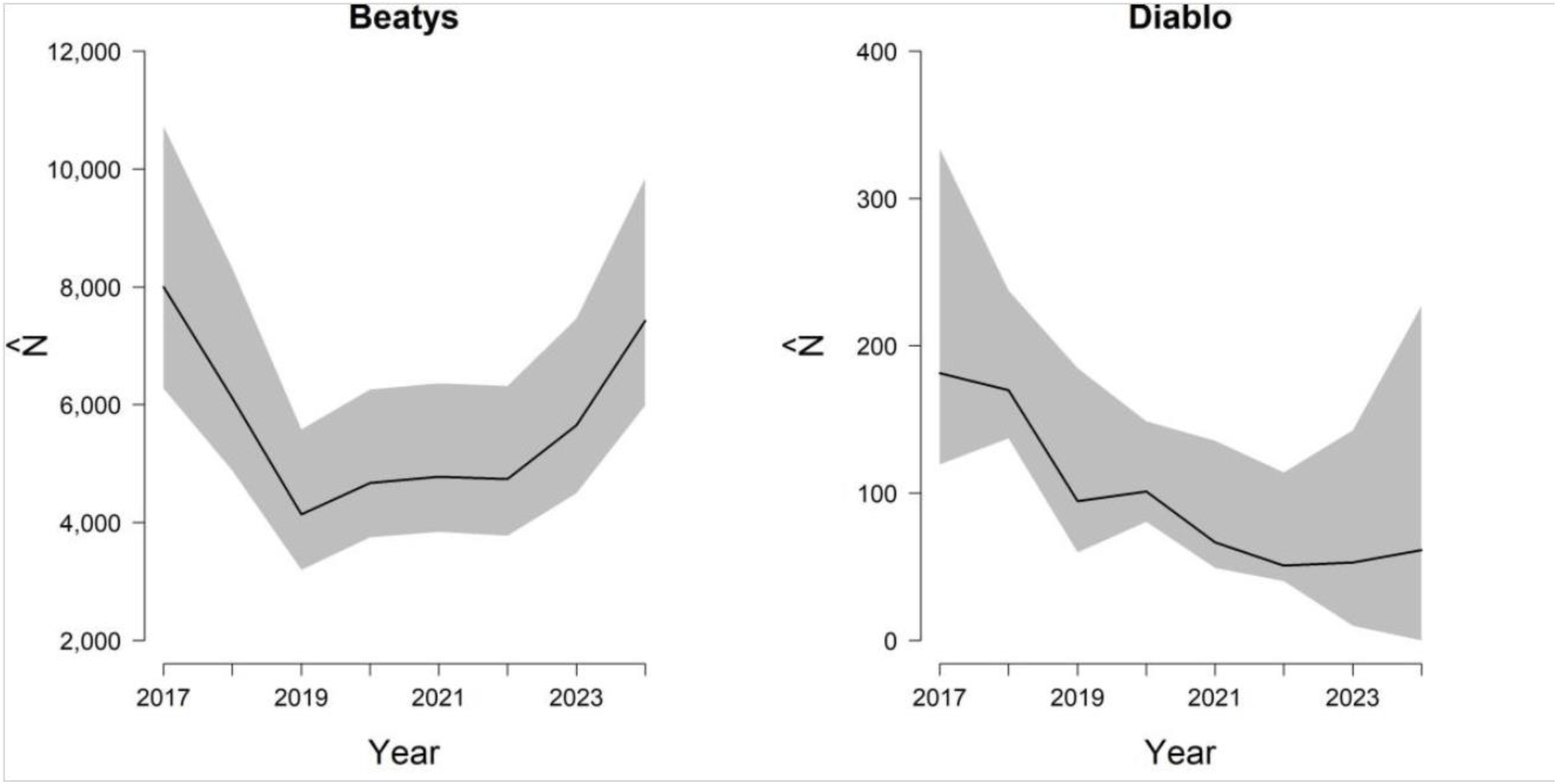
*N*-mixture model estimates of greater sage-grouse (*Centrocercus urophasianus*) abundance within Priority Areas for Conservation (PAC) located in the state of Oregon, USA between 2017–2024. PACs depicted (Beatys and Diablo) are identified in the title of each figure. The median (black line) and 95% credible interval (gray polygon) are plotted for each PAC.

The remaining 12 PACs cumulatively accounted for approximately 46.0% of the state’s total population size during 2024. The smallest of those PACs were Burns (151; 95% CRI = 117–237), Tucker Hill (215; 95% CRI = 171–320), and Saddle Butte (444; 95% CRI = 357–658), representing 1.9% of the 2024 total (Fig. 12).

**Figure 12.**
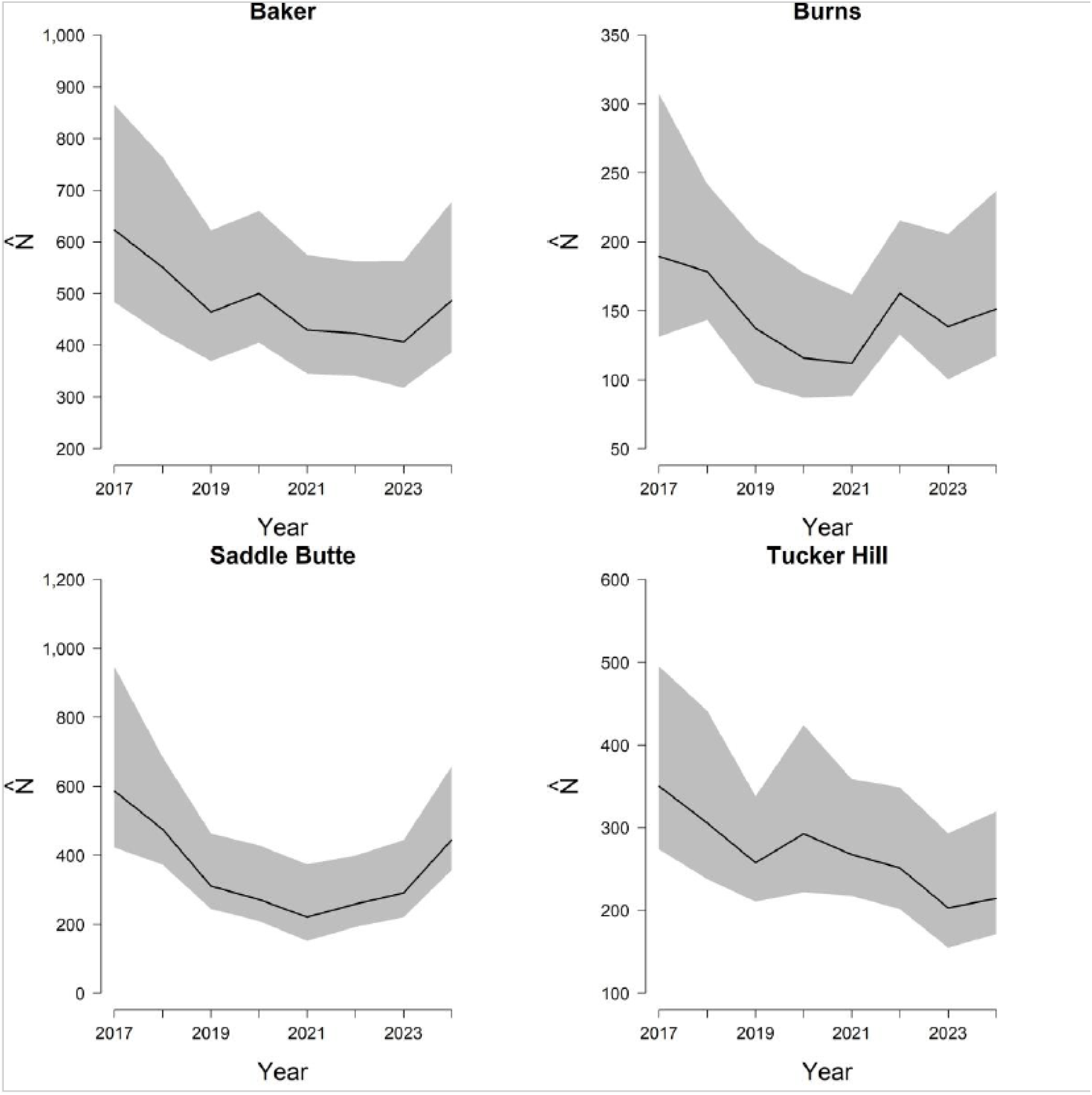
*N*-mixture model estimates of greater sage-grouse (*Centrocercus urophasianus*) abundance within Priority Areas for Conservation (PAC) located in the state of Oregon, USA between 2017–2024. PACs depicted (Baker, Burns, Saddle Butte, Tucker Hill) are identified in the title of each figure. The median (black line) and 95% credible interval (gray polygon) are plotted for each PAC.

**Figure 13.**
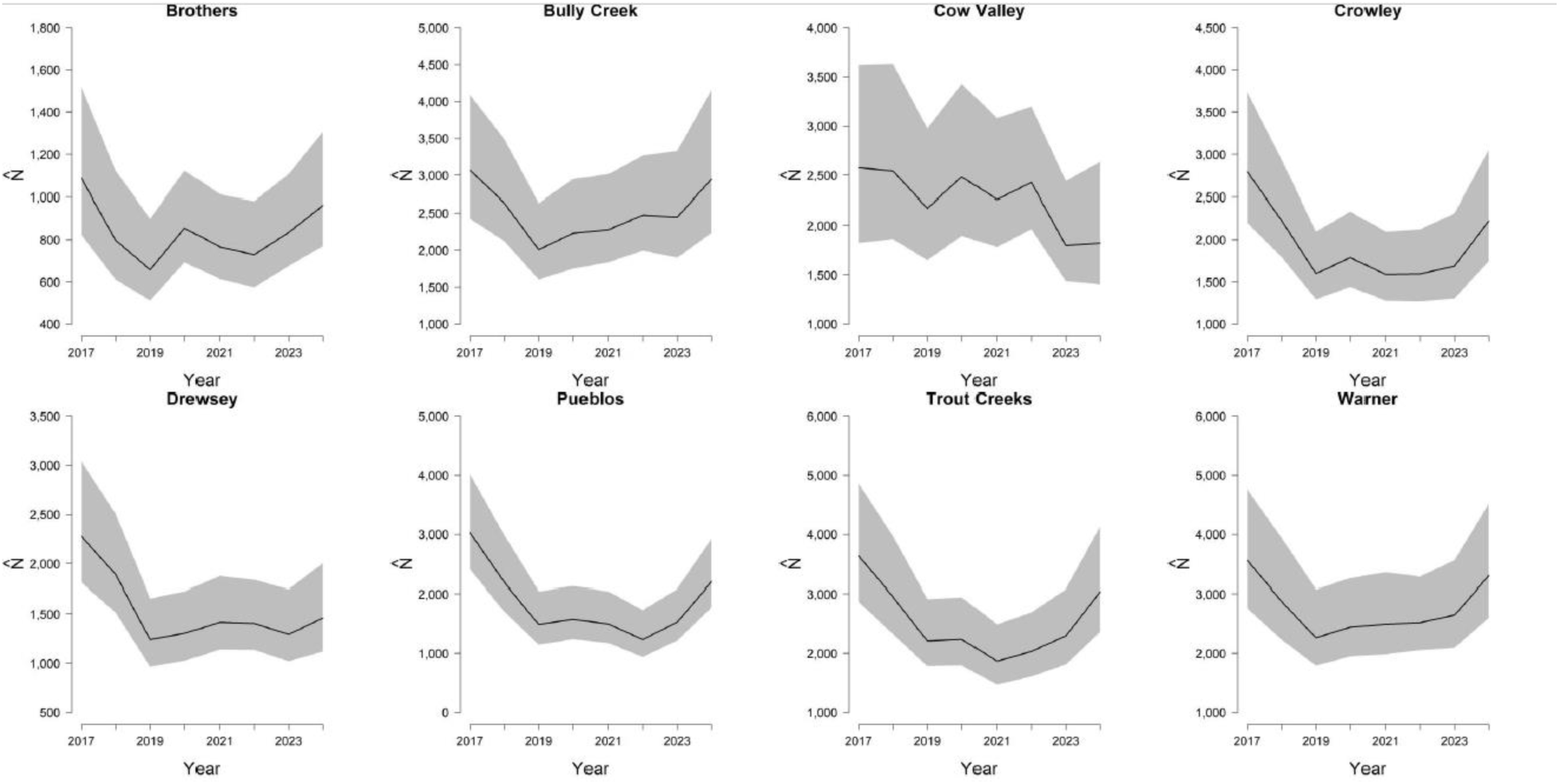
*N*-mixture model estimates of greater sage-grouse (*Centrocercus urophasianus*) abundance within Priority Areas for Conservation (PAC) located in the state of Oregon, USA between 2017–2024. PACs depicted (Brothers, Bully Creek, Cow Valley, Crowley, Drewsey, Pueblos, Trout Creeks, Warner) are identified in the title of each figure. The median (black line) and 95% credible interval (gray polygon) are plotted for each PAC.

Baker (487; 95% CRI = 386–678) and Brothers (961; 95% CRI = 769–1,307) contained considerably fewer sage-grouse compared to PACs of similar geographic size, which could be a product of their geographic location and general isolation (Figure 1). For example, the Cow Valley (1,821; 95% CRI = 1,402–2,637) and Bully Creek (2,947; 95% CRI = 2,234–4,154) PACs, located just south of Baker, share a common border with very little Low-Density areas between them (Figure 1). Additional border connections exist among Bully Creek, Crowley (2,221; 95% CRI = 1,745–3,052), and Drewsey (1,460; 95% CRI = 1,119–2,007), with each of those PACs having considerably larger population sizes compared to Baker. PACS possessing even greater connectivity potential (beyond the borders of Oregon), located in the southern portion of the state, include Pueblos (2,203; 95% CRI = 1,778–2,921), Trout Creeks (3,039; 95% CRI = 2,366–4,132), and Warner (3,301; 95% CRI = 2,604–4,517). Those PACs alone represented approximately 20% of the state’s total population size during 2024.

Low-density areas accounted for just 1.5% (634; 95% CRI = 445–965) of Oregon’s total population size during 2024 (Fig. 14) despite covering nearly 38% of the species’ mapped habitat within the state (Figure 1).

**Figure 14.**
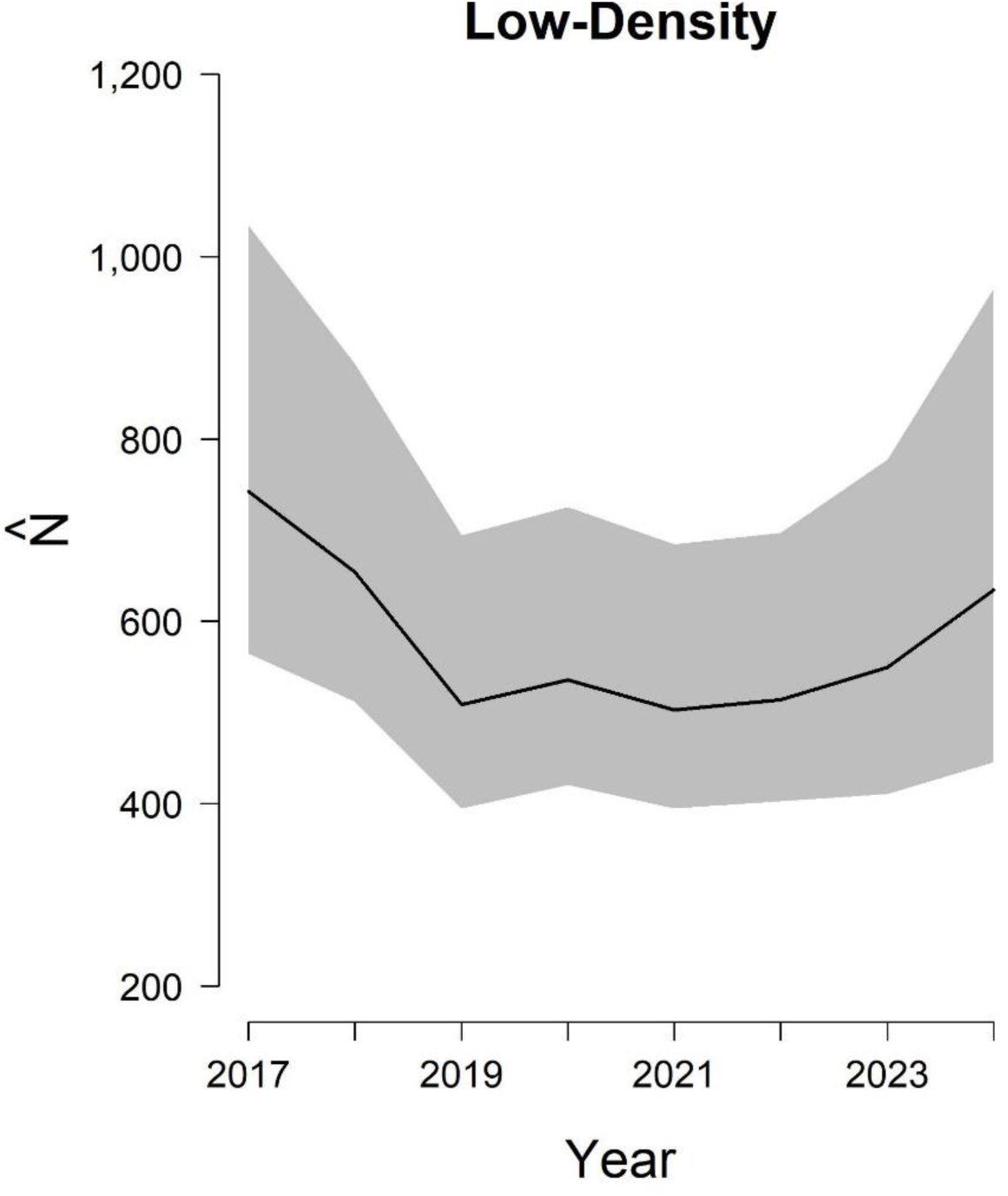
*N*-mixture model estimates of greater sage-grouse (*Centrocercus urophasianus*) abundance within Low-Density areas of the state of Oregon, USA between 2017–2024. The median (black line) and 95% credible interval (gray polygon) are plotted.

Evaluations of sampling intensity (percent of leks receiving repeated counts) revealed a negative trend between effort and bias with a reduced amount of effort corresponding to an increase in absolute bias (Table 2).

**Table 2.**
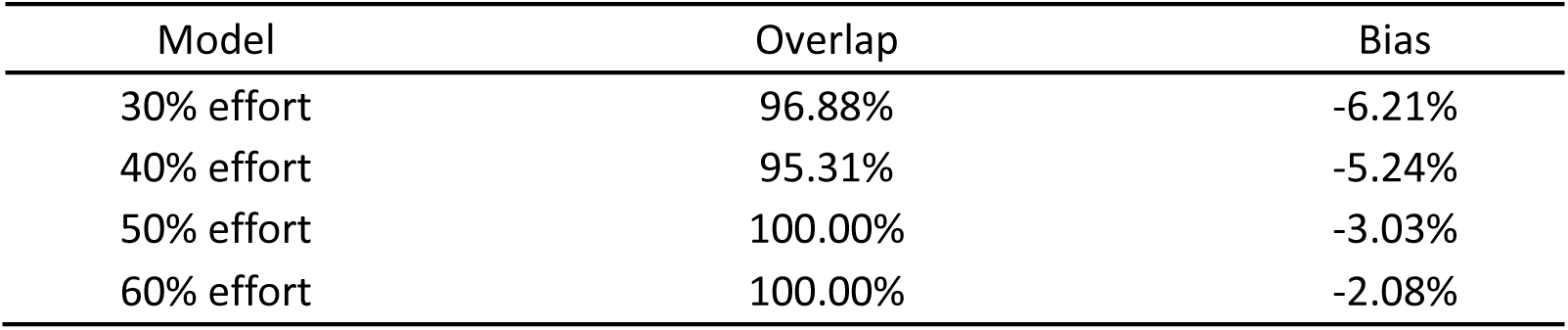
Evaluations of an N-mixture model fit to greater sage-grouse (*Centrocercus urophasianus*) lek (breeding area) count data collected within the state of Oregon, USA between 2017–2024. Four separate models with different levels of sampling intensity (30–60% of leks received repeated counts, *reduced models*) were compared to the same model fit to a dataset based on current maximal effort (70%, *reference model*). Bias was calculated as the mean difference between the median estimate of the *reference model* and the median estimate of the *reduced model*, divided by the median estimate of the *reference model*, multiplied by 100. Coverage was calculated as the percent of the *reduced model’s* 90% credible intervals (CRIs) that overlapped the median of the *reference model’s* counterpart parameters.

All bias estimates were negative indicating that as effort decreased, abundance estimates tended to increase. The average number of birds estimated per PAC between 2017–2024 was 1,560. Bias estimates of -2.08% (60% effort), -3.03% (50% effort), -5.24% (40% effort), and -6.21% (30% effort) corresponded to PAC-level abundance estimates that were 32, 46, 80, and 95 birds larger than the reference model, respectively. Using the 2024 median, state-wide estimate of 41,875 birds, those same percentages corresponded to abundance estimates that were 871 (60% effort), 1,269 (50% effort), 2,194 (40% effort), and 2,600 (30% effort) birds larger than the reference model. The increase in absolute bias was less than 1% (0.95%) between efforts equaling 60% and 50%, but more than doubled when reducing the effort from 50% to 40% (2.21%). The 90% CRIs overlapped the median estimate of the *reference model* 100% of the time for the 50% and 60% effort scenarios (Table 2), whereas 40% and 30% scenarios produced results that were less than 100% at 95.31% and 96.88%, respectively.

## Discussion

We fit an N-mixture model to within-season replicated lek counts of male sage-grouse collected from the state of Oregon, USA between 2017–2024. Models revealed a state-wide population estimate of 41,875 sage-grouse in 2024, which was down from a high of 50,869 in 2017 (start year of analysis). The population’s lowest point during the 8-year period occurred in 2019 when there was an estimated 30,644 sage-grouse across the entire state. Population management units (Priority Areas for Conservation; PACs) showed considerable variation in geographic size, making population comparisons difficult. However, PACs located in the south and southeastern portions of the state tended to support greater numbers and densities of birds. Moreover, they tended to exhibit greater post-nadir rebounds (i.e., increases in population size between 2019– 2024) despite comparable rates of bird loss leading up to the nadir (i.e., decreases in population size between 2017–2019). For example, the top four largest PACs, by 2024 population size (Beatys, Soldier Creek, Warner, Trout Creeks), declined 44% between 2017–2019 (20,034– 11,275), but increased 44% between 2019–2024 (11,275–16,251; net change = -18%).

Conversely, the four smallest PACs (Juniper Mountain, Sheepshead, Burns, Diablo) declined by a lesser amount (35%) between 2017–2019 (920–596) but made marginal gains (2%) after the nadir (596–612; net change = -34%). These initial results could signify the potential for an Allee effect and may suggest a need for future investigations to determine whether the smaller, isolated, and low-density PACs require greater attention or possible intervention (e.g., translocation).

The observation component of our model produced median lek-level detection probabilities that ranged from 0.23–0.82 (mean = 0.54). Investigations of temporal covariates provided some insights about the variability in detection, while being inconclusive for others. Specifically, the data and model supported a strong relationship between detection probability and time since sunrise with earlier times producing significantly higher rates of detection. Conversely, we failed to find support for a relationship between day of year and detection probability. We attribute this apparent lack of relationship to the narrow date range (11-March– 12-May) of counts used in this analysis, a period that corresponds with peak lek attendance and therefore lower variability in lek counts (Monroe et al., 2016; Wann et al., 2019). Based on these findings it may be possible to extend the sampling period slightly on both ends, particularly if the current window is constraining field operations associated with other projects or species. At the very least, these findings suggest that greater emphasis could be placed on the time of day rather than the day of year (restricted to the breeding season), when conducting a lek count within the state of Oregon.

We found that reductions in sampling effort corresponded with increases in absolute bias. Bias estimates were consistently negative with greater reductions in effort producing higher estimates of abundance. Specifically, we observed a 1.6% increase in absolute bias for every 10% reduction in effort. Though small, compared to the annual capacity of sage-grouse populations to grow and decline, these biases could lead to a significant amount of error when estimating population trends if observers introduced a strong temporal trend in sampling intensity (e.g., 10% reduction year over year for multiple years). This is something we did not observe for the data and period investigated. Despite the relationship to bias, we did not find a strong link between data sparsity and coverage (the percent of the *reduced model’s* CRIs that overlapped the median of the *reference model’s* parameters). Even the extremely disparate datasets (e.g., 30% effort) contained the target value within their CRIs more than 95% of the time. This would suggest minimal impact on long-term trend estimation. However, precision of estimates in year-over-year changes in population size could be drastically reduced. It is important to note that the simulated replacement of repeat counts with NA values was accomplished using a random process. Assuming managers implement a more targeted approach to reduced sampling in the future (e.g., narrow in on peak attendance hours and dates), the results could yield a less significant bias than what was estimated here.

## Data availability

The sage-grouse lek data used in this analysis are managed by the Oregon Department of Fish and Wildlife (ODFW). The ODFW can be contacted for information related to data availability.

## Acknowledgments

We conducted this analysis in close consultation with the Oregon Department of Fish and Wildlife (ODFW), which provided data and oversight. We thank Justin Small (Nevada Department of Fish and Wildlife) and Philip Gould (U.S. Geological Survey) for helpful comments in reviewing this manuscript in its entirety. We extend gratitude to Mikal Cline (ODFW) who provided feedback at various stages on uses of sage-grouse data, modeling methods, and constructive reviews at various stages of production. Any use of trade, firm, or product names is for descriptive purposes only and does not imply endorsement by the U.S. Government.

## Funding

This project could not have been completed without the financial support of the Oregon Department of Fish and Wildlife (ODFW) and U.S. Geological Survey.

## Conflict of interest disclosure

The authors of this preprint declare that they have no conflict of interest relating to the content of this article.

## Notes

### Competing Interest Statement

The authors have declared no competing interest.

